# Early survival and delayed death of developmentally-born dentate gyrus neurons

**DOI:** 10.1101/092957

**Authors:** Shaina P. Cahill, Ru Qi Yu, Dylan Green, Evgenia V. Todorova, Jason S. Snyder

## Abstract

The storage and persistence of memories depends on plasticity in the hippocampus. Adult neurogenesis produces new neurons that mature through critical periods for plasticity and cellular survival, which determine their contributions to learning and memory. However, most granule neurons are generated prior to adulthood; the maturational timecourse of these neurons is poorly understood compared to adult-born neurons, but is essential to identify how the dentate gyrus, as a whole, contributes to behavior. To characterize neurons born in the early postnatal period, we labeled dentate gyrus neurons born on postnatal day 6 (P6) with BrdU and quantified maturation and survival across early (1 hour to 8 weeks old) and late (2-6 months old) cell ages. We find that the dynamics of developmentally-born neuron survival is essentially the opposite of neurons born in adulthood: P6-born neurons did not go through a period of cell death during their immature stages (from 1-8 weeks). In contrast, 17% of P6-born neurons died after reaching maturity, between 2-6 months of age. Delayed death was evident from the loss of BrdU^+^ cells as well as pyknotic BrdU^+^caspase3^+^ neurons within the superficial granule cell layer. Patterns of DCX, NeuN and activity-dependent Fos expression indicate that developmentally-born neurons mature over several weeks and a sharp peak in zif268 expression at 2 weeks suggests that developmentally-born neurons mature faster than adult-born neurons (which peak at 3 weeks). Collectively, our findings are relevant for understanding how developmentally-born dentate gyrus neurons contribute to memory and disorders throughout the lifespan. High levels of early survival and zif268 expression may promote learning, while also rendering neurons sensitive to insults at defined stages. Late neuronal death in young adulthood may result in the loss of hundreds of thousands of dentate gyrus neurons, which could impact memory persistence and contribute to hippocampal/dentate gyrus atrophy in disorders such as depression.

## INTRODUCTION

Many hippocampal dentate gyrus (DG) granule neurons are born postnatally in rodents (Crespo et al., 1986; Cameron and McKay, 2001; Rao and Shetty, 2004). The adult-born population alone ultimately comprises up to 40% of the DG population in rats (Snyder and Cameron, 2012a), and a large number in mice as well (DeCarolis et al., 2013). Recent data suggests that cumulative neurogenesis, over decades, may produce similarly large proportions of DG neurons in adult humans (Spalding et al., 2013). Despite the substantial number of neurons added in adulthood, many DG neurons are generated during the perinatal period and yet comparably little is known about their properties. A better characterization of developmentally-born neurons is necessary to understand how different populations of neurons, born at different periods of life, contribute to DG function as a whole.

The basic pattern of DG development has been described. Precursor cells migrate from the dentate notch in the late embryonic period and form a germinal zone in the hilus that gives rise to granule neurons (Altman and Bayer, 1990a; 1990b). In the rat, 1-10% of DG neurons are added daily between E14 and P14 (Schlessinger et al., 1975). Thus, at any given time in the first few weeks of life the DG is made up of a heterogeneous population of neurons that span all stages of cellular development. This heterogeneity is reflected in patterns of electrophysiology, morphology and immediate-early gene (IEG) expression (Liu et al., 2000; Jones et al., 2003; Montes-Rodríguez et al., 2013). However, in the absence of cellular birthdating methods, it is impossible to identify how anatomical and functional properties relate to neuronal age.

Markers of dividing cells, such as tritiated thymidine, BrdU and retroviruses, have been used extensively to characterize the birth, survival and cellular properties of DG neurons, particularly those born in adulthood. These studies have revealed that adult-born neurons go through a sensitive period of 4 weeks during which many neurons die (Cameron et al., 1993; Kempermann et al., 2003; McDonald and Wojtowicz, 2005; Tashiro et al., 2006; Mandyam et al., 2007; Snyder et al., 2009a), after which neurons survive indefinitely (Dayer et al., 2003; Kempermann et al., 2003). During immature stages adult-born neurons form afferent and efferent synapses according to a specific pattern (Espósito et al., 2005), they display critical periods for synaptic plasticity and memory (Ge et al., 2007; Gu et al., 2012), and their survival can be modulated by learning, stress, exercise and enriched environment, often at very specific cell ages (Döbrössy et al., 2003; Olariu et al., 2005; Dupret et al., 2007; Epp et al., 2007; Tashiro et al., 2007; Snyder et al., 2009b; Alvarez et al., 2016).

In contrast to adult neurogenesis, such extensive and detailed analyses of developmental DG neurogenesis are lacking. Studies that have employed birthdating methods indicate that developmentally-born neurons mature faster than adult-born neurons (Overstreet-Wadiche et al., 2006; Zhao et al., 2006). Electrophysiological properties may ultimately be similar once cells have reached maturity (Laplagne et al., 2006). However, different immediate-early gene responses to experience (Tronel et al., 2014) and enhanced morphological plasticity in old adult-born cells (Tronel et al., 2010) suggest that there may be persistent differences between DG neurons born at different ages. Understanding the behavioral contribution of developmentally-born neurons also depends on their patterns of survival. For example, neuronal survival may contribute to persistent information storage and yet there is no detailed quantification of the initial lifespan of developmentally-born neurons, though there is evidence that they may die in adulthood (Dayer et al., 2003).

To better understand the maturation and survival of developmentally-born DG neurons we used the thymidine analog BrdU to label DG neurons born at postnatal day 6 (P6). We found that, unlike neurons born in adulthood, P6-born neurons do not undergo appreciable cell death during their immature stages but rather undergo delayed cell death between 2-6 months of age. Furthermore, patterns of immediate-early gene expression suggest P6-born neurons mature faster than adult-born neurons and show peak zif268 expression when they are 2 weeks old. Collectively, these unique patterns of neuronal maturation and turnover suggest that developmentally-born neurons may contribute unique forms of plasticity to the DG throughout the lifespan, which could play a role in the dynamic nature of hippocampal memory.

## METHODS

### Animals and treatments

All procedures were approved by the Animal Care Committee at the University of British Columbia and conducted in accordance with the Canadian Council on Animal Care guidelines regarding humane and ethical treatment of animals. Experimental Long-Evans rats were generated in the Department of Psychology animal facility with a 12-hour light/dark schedule and lights on at 6:00 am. Breeding occurred in large polyurethane cages (47 cm × 37 cm × 21 cm) containing a polycarbonate tube, aspen chip bedding and ad libitum rat chow and water. The day of birth was designated postnatal day 1. Litters ranged from 8-18 pups, and pups from each litter were distributed equally amongst experimental groups. Breeders (both male and female) remained with the litters until P21, when offspring were weaned to 2 per cage in smaller polyurethane bins (48 cm × 27 cm × 20 cm).

This study is comprised of 2 experiments that examine developmentally-born neurons at early vs. late intervals. In both experiments rats were injected with the thymidine analog BrdU (50 mg/kg, intraperitoneal) at P6, to label neurons born at the peak of granule cell birth (Schlessinger et al., 1975). In experiment 1, to track early neuronal survival and development, equal numbers of male and female rats were killed at the following post-BrdU injection timepoints: 1 hour, 1 day, 3 days, 1 week, 2 weeks, 3 weeks, 4 weeks, 8 weeks. One hour before being euthanized, rats in the 1-8 week groups were exposed to a novel environment to induce activity-dependent IEG expression. The novel environment exposure consisted of 1 hour in an empty cage filled with corncob bedding in an unfamiliar room. Rats were picked up and briefly handled at least twice during the exposure. Immediately after the novel environment exposure rats were anaesthetized with isoflurane and perfused with 4% paraformaldehyde in phosphate buffered saline (PBS, pH 7.4). Brains remained in paraformaldehyde for 48 hours and were then stored in 0.1 % sodium azide in PBS until processed. In experiment 2, to examine cell death and the long-term survival of cells born on P6, male rats were killed at 2 months and 6 months of age (when BrdU^+^ cells were 8 weeks old and 26 weeks old, respectively; for clarity we subsequently refer to these timepoints as 2 and 6 months). These rats did not receive any environmental exposure for immediate-early gene analyses, but were perfused directly from their home cage.

### Tissue processing and immunohistochemistry

Brains were immersed in 10% glycerol solution for 1 day, 20% glycerol solution for 2 days and then sectioned coronally at 40 *μ*m on a freezing microtome. Sections were stored in cryoprotectant at 20°C until immunohistochemical processing. For stereological quantification of BrdU^+^ cells a 1 in 12 series of sections throughout the entire dentate gyrus were mounted onto slides, heated to 90°C in citric acid (0.1 M, pH 6.0), permeabilized in PBS with 10% triton-x for 30min and then incubated overnight with goat anti-Prox1 (1:1000 in 10% triton-x and 3% horse serum; R&D systems, AF2727). Sections were washed, incubated in biotinylated donkey anti-goat secondary antibody for 1 hour (1:250; Jackson, 705-065-147), 0.3% hydrogen peroxide for 30 min and Prox1^+^ cells were visualized with an avidin-biotin-horseradish peroxidase kit (Vector Laboratories) and vector nova red HRP substrate (Vector Laboratories). Sections were washed, permeabilized with trypsin, incubated in 2N HCl for 30 min to denature DNA, and incubated overnight with mouse anti-BrdU (1:200 in 10% triton-x and 3% horse serum, BD Biosciences; 347580). Sections were washed and incubated in biotinylated goat anti-mouse secondary antibody for 1 hour (1:250; Sigma, B0529), and BrdU^+^ cells were visualized with an avidin-biotin-horseradish peroxidase kit (Vector Laboratories) and cobalt-enhanced DAB (Sigma Fast Tablets). Sections were then rinsed in PBS, dehydrated, cleared with citrisolv (Fisher) and coverslipped with Permount (Fisher).

Immunohistochemical analyses of neuronal phenotype were performed on free-floating sections with fluorescent detection. Sections were treated with 2N HCl for 30 minutes, incubated at 4°C for 3 days in PBS with 10% triton-x, 3% horse serum and combinations of the following antibodies: rat anti-BrdU (1:200; AbD Serotec, OBT0030G), goat anti-doublecortin (1:250; Santa Cruz, sc-8066), mouse anti-NeuN (1:200; Millipore, MAB377), rabbit anti-zif268 (1:1000; Santa Cruz, sc-189), goat anti-c-fos (1:250; Santa Cruz, sc-52G), rabbit anti-active caspase3 (1:200 BD Biosciences, 559565) and mouse anti-PCNA (1:200 Santa Cruz, sc-56). Visualization was performed with Alexa488/555/647-conjugated donkey secondary antibodies (Invitrogen/Thermofisher) diluted 1:250 in PBS for 60 minutes at room temperature. Sections were counterstained with DAPI, mounted onto slides and coverslipped with PVA-DABCO.

### Microscopy and sampling

Quantification of total DAB-stained BrdU^+^ cells was performed under brightfield microscopy using stereological principles. A 1 in 12 series of sections spanning the entire dentate gyrus was examined with a 40x objective and an Olympus CX41 microscope. All BrdU^+^ cells located within the granule cell layer or its 20 *μ*m hilar border (the subgranular zone) were counted in each section and counts were multiplied by 12 to estimate the total number of BrdU^+^ cells per DG (bilaterally). At the 1hr, 1d and 3d timepoints (postnatal days 6-9) the DG was not yet fully formed and many BrdU^+^ cells resided in the tertiary dentate matrix, located in the hilus (Altman and Bayer, 1990a). To accurately capture BrdU^+^ cells from all regions that will ultimately form the DG granule cell layer we used the dentate granule neuron-specific marker Prox1 to delineate the boundary of the DG and its germinal zones and counted all BrdU^+^ cells in this region (Fig. 1A).

**Figure 1:**
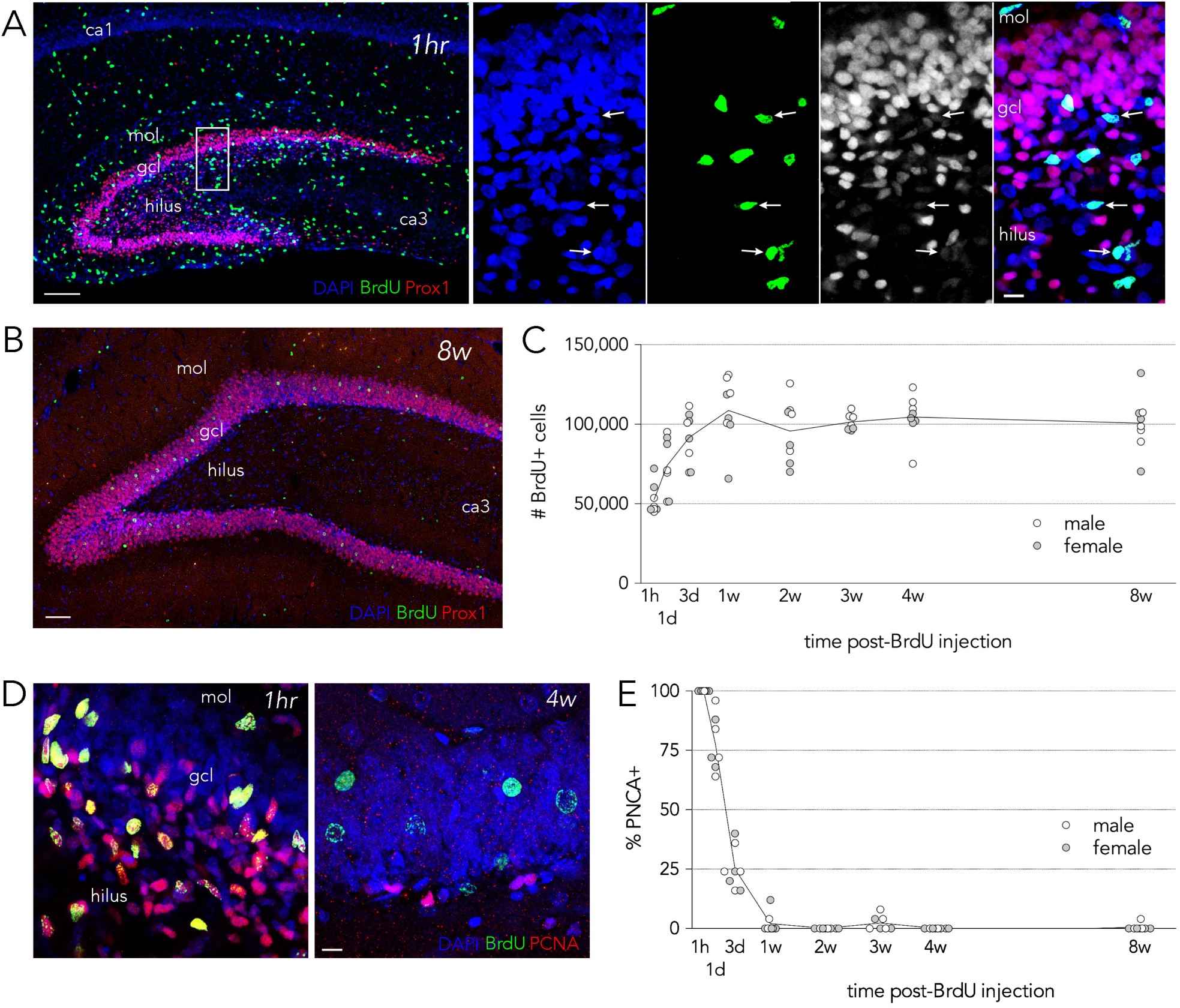
Dynamics of developmental neurogenesis and early cell survival. ***A***) Confocal images of the dentate gyrus 1 hour after BrdU injection at P6. At this early stage of dentate gyrus formation many BrdU^+^ cells can be found within the granule cell layer (gcl) and the proliferative tertiary dentate matrix in the hilus. Granule neuron precursors and granule neurons can be identified by immunoreactivity for the granule neuron-specific marker Prox1. Inset shows BrdU^+^ and weakly Prox1^+^ cells in the granule cell layer and hilus (arrows). Scale bars, 100 *μ*m and 10 *μ*m for the low and high magnification images, respectively. ***B***) Confocal image of the dentate gyrus 8 weeks after P6 BrdU injection. BrdU^+^ granule neurons are limited to the granule cell layer. Scale bar, 100 *μ*m. ***C***) The total number of BrdU^+^ cells labeled at P6 doubled from 1 hour (52,407) to 1 week (108,593) and remained stable thereafter (n = 7-8/timepoint; ANOVA F_7,54_ = 10.8, P < 0.0001). BrdU^+^ cell number was significantly greater at 3 days and 1 week compared to 1 hour (Ps < 0.001). BrdU^+^ cell number did not differ between any of the 3 day to 8 week timepoints (all Ps > 0.4). ***D***) Confocal images of the dentate gyrus immunostained for BrdU and the proliferation marker PCNA reveal many double labeled cells at 1 hour but not 4 weeks. Scale bar, 10 *μ*m. ***E***) There was a significant decline in BrdU^+^ cells that expressed PCNA, where all BrdU^+^ cells expressed PCNA at 1 hour, 78% at 1 day and 25% at 3 days (ANOVA F_7,54_ = 429, P < 0.0001; post hoc comparisons all P < 0.0001). Data for males and females are indicated in the graphs but were pooled for statistical analyses. Mol, molecular layer; gcl, granule cell layer.

Fluorescent tissue was examined on a confocal microscope (Leica SP8). For PCNA, DCX and NeuN approximately 25 BrdU^+^ cells per animal located in the suprapyramidal blade of the dorsal DG were examined for marker co-expression using a 63X objective (NA 1.4) and 1 *μ*m z-sections throughout each cell. Activity in P6-born neurons was assessed by examining the immediate-early genes Fos and zif268 in 200 BrdU^+^ cells per animal, also sampled from the suprapyramidal blade of the dorsal DG, using a 40X oil-immersion lens (NA 1.3) and offline analyses of image stacks. Since zif268 and Fos staining intensity in immature neurons is graded, fluorescence intensity for each cell was measured and compared to background levels (regions in the hilus devoid of DAPI^+^ cell bodies) within each field. Cells were counted as positive if staining intensity was twice background, a threshold that captures cells with at least moderate levels of immediate-early gene immunostaining (Snyder et al., 2009b). Activity in the overall population of DG neurons was assessed by examining Fos and zif268 expression in ~300 DAPI^+^ granule neurons from the same image stacks as the BrdU analyses. Approximately equal numbers of DAPI^+^ cells were sampled from two regions of the suprapyramidal blade, located 1/3 and 2/3 along its medial-lateral extent, and spanning layers of the granule cell layer (to ensure cells of all ages were sampled equally). As with zif268 and Fos, NeuN expression increases will cell age/maturity (Snyder et al., 2009a). We therefore quantified both weak NeuN expression (2x background, to quantify all neurons) and strong NeuN expression (4x, to quantify relatively mature neurons; see Fig. 2B for examples). For cell death measurements in 2 and 6-month-old rats we found that expression of caspase3 was widespread, graded and only cells that strongly expressed caspase3 were also pyknotic. To avoid overestimating numbers of dying cells we therefore only quantified cells that had intense immunostaining for caspase3 and were also pyknotic. A 1 in 12 series of sections spanning the entire dentate gyrus was examined for dying cells and dying BrdU^+^ cells. Immature neurons tend to reside in the deep layers near the hilus/subgranular zone and older neurons reside in the superficial layers, near the molecular layer (Crespo et al., 1986; Wang et al., 2000; Muramatsu et al., 2007; Mathews et al., 2010). To estimate the age of dying cells, each pyknotic caspase3^+^ cell was characterized according to its anatomical position within the granule cell layer. The thickness of the granule cell layer was normalized to 100, with the hilar border being 0 and the molecular layer border being 100. The position of dying cells was measured as the relative distance from the hilar border to the middle of the cell body. Cells located in the subgranular zone were assigned a score of 0. To compare the anatomical distribution of dying cells with the distribution of developmentally-born and adult-born cells, we analyzed the distribution of 25 P6-labeled BrdU^+^ cells and 25 DCX^+^ cells from each rat.

**Figure 2:**
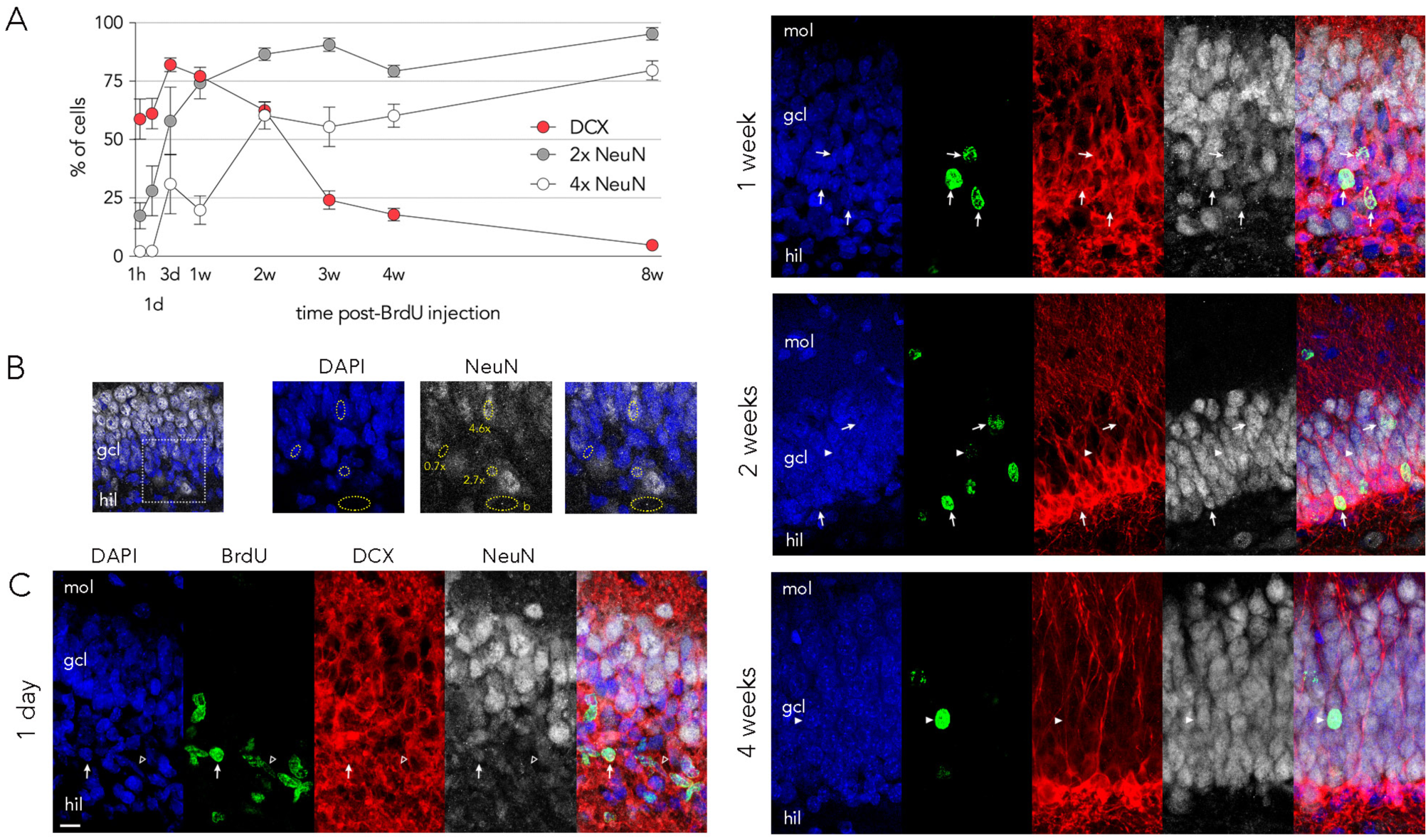
Early timecourse of neuronal marker expression. ***A***) The vast majority of BrdU^+^ cells were neuronal from 3 days (82% DCX^+^) until 8 weeks (95% 2x background NeuN^+^). Just over half of BrdU^+^ cells expressed DCX at the 1 hour and 1 day timepoints before reaching maximal expression at 3 days (ANOVA F_7,52_ = 38, P < 0.0001; 1d vs 3d P = 0.04, 3d vs 2w P = 0.047, 2w vs 3w P < 0.0001, 2w vs 8w P = 0.06). 2x NeuN expression increased rapidly during the first week and then stabilized (ANOVA F_7,52_ = 17, P < 0.0001; 1h and 1d vs 1w Ps < 0.001, differences between 1w-8w timepoints all P > 0.4). Stronger 4x NeuN expression increased markedly over the first two weeks and continued to rise to 80% by 8 weeks (ANOVA F_7,52_ = 20, P < 0.0001; 1d vs 3d P = 0.04, 1w vs 2w P = 0.0005, differences between 2-8w timepoints all P > 0.16). ***B***) Raw confocal images (single z-plane) illustrating NeuN expression-levels. DG neuron ROIs and corresponding NeuN intensities relative to background (“b”) are indicated. ***C***) Confocal images of P6-born neurons immunostained for BrdU, DCX and NeuN. <u>One day</u> after BrdU injection, on P7, the granule cell layer was dispersed and BrdU^+^ cells were scattered throughout the hilus bordering the granule cell layer. More mature granule neurons, in the superficial layers near the molecular layer, had stronger NeuN immunoreactivity. The entirety of the dentate gyrus showed DCX^+^ immunoreactivity. The arrow indicates a BrdU^+^ cell that expresses DCX and NeuN. The open arrowhead indicates a BrdU^+^ cell that did not stain for NeuN. <u>One week</u> after BrdU injection (P13) there was a strong gradient of NeuN expression across cell layers and most BrdU^+^ cells expressed both DCX and NeuN (arrows). <u>Two weeks</u> after BrdU injection some BrdU^+^ cells expressed both DCX and NeuN (arrows) and some only expressed NeuN (arrowhead). At <u>four weeks</u> post-BrdU injection the granule cell layer was adult-like, with a thin, tight layer of DCX^+^ cells bordering the hilus and no gradient of NeuN expression; BrdU cells overwhelmingly expressed NeuN and not DCX (see arrowhead). Mol, molecular layer; gcl, granule cell layer; hil, hilus.

### Statistical analyses

Developmental timecourse analyses (1hr to 8w groups) were performed using ANOVA with Holm-Sidak post hoc tests corrected for multiple comparisons. Differences between the long-term survival groups (2mos and 6mos) were assessed by unpaired t test or Mann Whitney test. In all cases significance was set at p = 0.05.

## RESULTS

### Early survival of developmentally-born DG neurons

A single BrdU injection at P6 labeled many DG neurons. At early timepoints (1hr, 1d and 3d post-injection BrdU^+^ cells could be observed throughout the proliferative tertiary dentate matrix in the hilus and in the deep layers of the granule cell layer (Fig. 1). At longer survival intervals BrdU^+^ cells were found throughout the granule cell layer but most were observed in the middle layers. Quantitatively, there was a steady increase in BrdU+ cells from 1hr to 1w, resulting in a doubling of cells from ~50,000 cells to just over 100,000 cells (Fig. 1C). Co-labeling with PCNA, a marker of cell division, revealed that 100%, 78% and 25% of BrdU^+^ neurons were undergoing cell division at the 1hr, 1d and 3d timepoints, respectively (Fig. 1D, E). Thus, growth of the BrdU^+^ cell population is due to the continued division of BrdU-labeled precursor cells until BrdU is diluted beyond the limits of detection, as has been observed in adult animals (Dayer et al., 2003). At 1 week and beyond, a negligible proportion of BrdU^+^ cells were labeled with PCNA. It is well established that adult-born neurons transition through a critical period for survival, when neurons are between 1 and 4 weeks of age and many undergo cell death (Cameron et al., 1993; Gould et al., 1999; Brandt et al., 2003; Snyder et al., 2009a). In contrast to the adult pattern, we found that BrdU^+^ cell numbers remained constant from 1-8 weeks of age, suggesting developmentally-born neurons do not undergo appreciable cell death during this period (and may be culled primarily at earlier stages of cellular development (Gould et al., 1994)). We observed no obvious differences in developmental neurogenesis between males and females but our small sample precludes any definitive conclusions (n=3-4/sex/timepoint).

### Expression of doublecortin and NeuN

To assess the rate of maturation of P6-born DG neurons we first quantified expression of the immature neuronal marker DCX and the mature neuronal marker NeuN, both of which have been extensively characterized in adult-born neurons. At the 1hr to 3d timepoints DCX was expressed at very high levels throughout the granule cell layer and the tertiary dentate matrix in the hilus (Fig. 2). Given the density of DCX^+^ processes it was difficult to unambiguously characterize all neurons as either DCX positive or negative. However, many cells were surrounded by DCX immunoreactivity and even expressed weak levels of NeuN at these early timepoints. Approximately ~60% of BrdU^+^ cells expressed DCX at the 1hr and 1d timepoints, when the majority of cells were also PCNA^+^. Thus, as in the adult brain (Brown et al., 2003), a substantial proportion of actively dividing precursor cells are committed to a neuronal lineage in the developing brain. The proportion of BrdU^+^ cells that were DCX^+^ rose to ~80% at both 3d and 1w as daughter cells matured into a post-mitotic neuronal phenotype. Thereafter, DCX expression waned but remained at 20-25% at 3-4 weeks. As previously observed in adult-born cells, NeuN expression increased with cell age and distinct patterns were observed for weak vs. strong levels of NeuN immunoreactivity (Brown et al., 2003; Snyder et al., 2009a). Whereas strong NeuN expression was not observed at the earliest timepoints, and increased over several weeks, weak NeuN expression was observed as early as 1 hour post-BrdU injection and reached near-peak levels by 2 weeks.

### Expression of activity-dependent immediate-early genes

Immediate-early gene expression is upregulated by synaptic activity and is often used as a proxy for neuronal activity, to identify experience-dependent recruitment of neuronal ensembles (Worley et al., 1991; Guzowski et al., 1999; Satvat et al., 2011). In adult-born neurons, expression of genes such as zif268, Fos and Arc develops over several weeks as new neurons integrate into circuits (Jessberger and Kempermann, 2003; Kee et al., 2007; Snyder et al., 2009a). To identify the rate at which P6-born neurons are integrated into circuits and recruited during exploration of a novel environment, we quantified zif268 and Fos expression at the 1, 2, 3, 4 and 8 week timepoints (Fig. 3). We found that zif268 was only expressed in 2% of 1-week-old BrdU^+^ cells but there was a dramatic but transient peak in expression at the 2 week timepoint, to 16% (Fig. 3B). Expression then dropped to 7%, 10%, and 12% at 3, 4 and 8 weeks, respectively. Notably, while P6-born cells had transient peak expression at 2 weeks, this pattern was not observed in the overall population of DG neurons, which gradually increased zif268 expression from 4% at the 1 week timepoint to 9% at the 4 week timepoint (corresponding to animal ages of ~2 to 5 weeks). In contrast to zif268, Fos expression was much lower and increased to ~1% of BrdU^+^ cells by the 4 week timepoint (Fig. 3C). Fos expression in the general DG population followed a similar pattern, though expression levels were significantly greater than in BrdU^+^ cells, possibly because the overall population of DG neurons was, on average, older than BrdU^+^ cells. To investigate whether zif268 and Fos identified different populations of cells activated by experience we quantified the amount of overlapping immediate-early gene expression in BrdU^+^ cells (Fig. 3D). Since both zif268 and (especially) Fos are expressed in only a fraction of cells, counts were pooled across animals within each timepoint. Generally, BrdU^+^ cells that expressed Fos also expressed zif268 (86% of cells across 2-8w timepoints, range 67-100%). In contrast, only 4% of BrdU^+^zif268^+^ cells also expressed Fos (range 1-8% across 2-8w timepoints). This was driven by the large number of 2-week-old BrdU^+^zif268^+^ neurons that did not express Fos (only 2/266 co-expressed Fos, < 1%). Older BrdU^+^zif268^+^ cells were more likely to also express Fos but the pattern of partial overlap suggests that zif268 may identify a population of active neurons that are not captured by Fos immunostaining.

**Figure 3:**
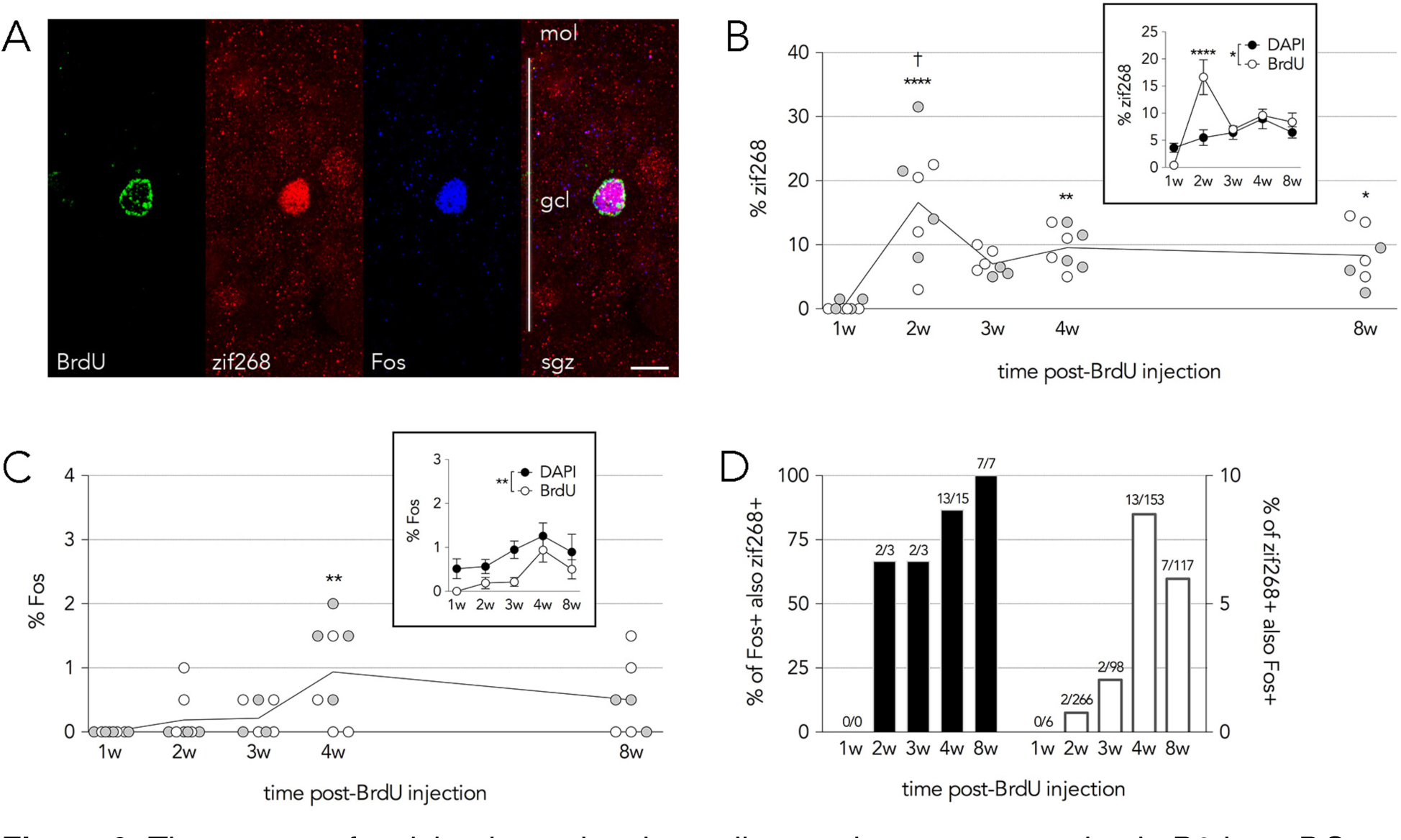
Timecourse of activity-dependent immediate-early gene expression in P6-born DG neurons and the general population of DG neurons. ***A***) Confocal image of an 8-week-old BrdU^+^ dentate gyrus neuron expressing zif268 and Fos following exploration of a novel environment. Vertical white line indicates the width of the granule cell layer (gcl). Scale bar = 10 *μ*m. mol, molecular layer; sgz, subgranular zone. ***B***) zif268 expression was virtually absent in 1-week-old cells, peaked at 2 weeks (16%), and then stabilized at 3-8 weeks (7-10%; ANOVA F_4,33_ = 11, P < 0.0001). ^†^zif268 expression at 2 weeks was greater than all other time points (all Ps < 0.05) and expression levels at 3, 4 and 8 weeks were not different from each other (Ps > 0.6). ****P < 0.0001, **P < 0.01, * P < 0.05 vs 1 week. Inset: Comparison of zif268 expression in P6-born BrdU^+^ cells and DAPI^+^ cells (overall population). zif268 expression was greater in BrdU^+^ cells than in DAPI^+^ cells at the 2 week timepoint (2 way repeated measures ANOVA effect of time F_4,32_ = 8, P = 0.0002, effect of cell population F_1,32_ = 5, *P < 0.05, interaction F_4,32_ = 7, P = 0.0005; ****P < 0.0001). ***C***) Fos expression in BrdU^+^ cells gradually increased, reaching 1% by 4 weeks (ANOVA F_4,33_ = 5, P < 0.01), and occurred at much lower rates than zif268. Fos expression at 4 weeks, but not the other timepoints, was significantly greater than at 1 week (**P < 0.01). Inset: Fos expression increased over time and was greater in the general population of DG neurons (DAPI^+^ cells) than in P6-born BrdU^+^ cells. Fos expression was not significantly different between BrdU^+^ cells and DAPI^+^ cells at any timepoint (2 way repeated measures ANOVA, effect of cell population F_1,32_ = 10, P = 0.004; effect of time F_4,32_ = 4, P = 0.006; interaction F_4,32_ = 0.2, P = 0.9). ***D***) Co-expression of zif268 and Fos in BrdU^+^ cells. Most BrdU^+^Fos^+^ cells also expressed zif268 but few BrdU^+^zif268^+^ cells also expressed Fos. Numbers above each bar indicate the fraction of cells expressing both immediate-early genes (pooled across animals). Sexes are pooled for all statistical analyses; lines indicate group means; error bars indicate s.e.m.

### Late death of developmentally-born cells

The early timecourse data indicate that P6-born neurons are remarkably stable during their immature stages, unlike adult-born neurons. Adult-born neurons do not die after reaching maturity (once 4 weeks old)(Dayer et al., 2003; Kempermann et al., 2003) but less is known about the long-term survival of developmentally-born neurons. We therefore injected male rats with BrdU at P6 and compared the number of BrdU^+^ cells at 2 and 6 months of age. In contrast to the initial stability of P6-born neurons, we found a significant loss of BrdU^+^ cells (17% or 29,000 cells) over this 4 month period of early adulthood (Fig. 4B).

**Figure 4:**
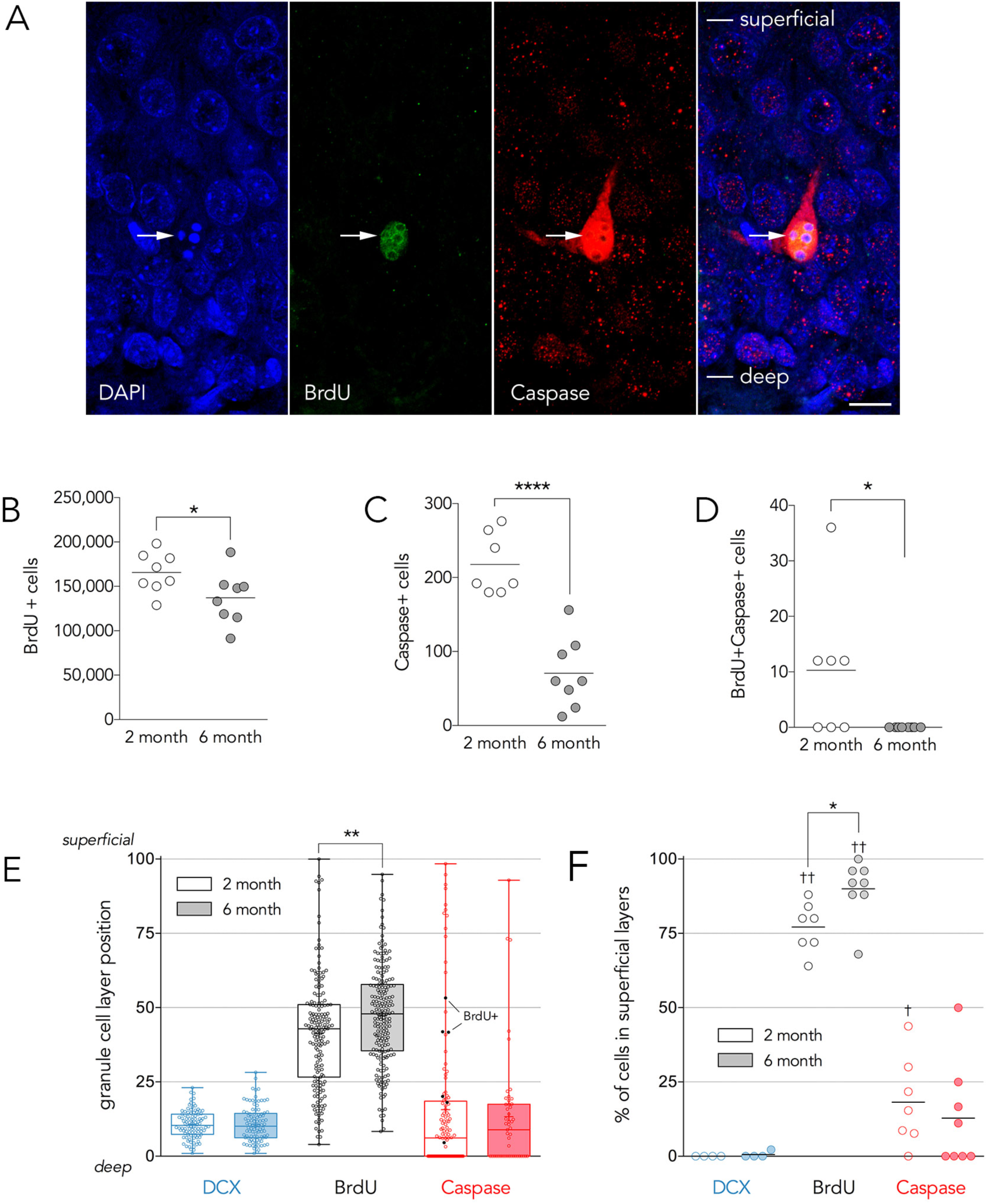
Delayed death of developmentally-born neurons. ***A***) Confocal image of a pyknotic caspase3^+^BrdU^+^ cell, undergoing apoptosis (arrow). Lines indicate the superficial and deep borders of the granule cell layer that were used to calculate the anatomical distribution of dying dentate gyrus neurons. ***B***) Between 2 and 6 months of age there was a 17% loss of BrdU^+^ cells that were born on P6 (166,000 cells at 2 months, 137,000 cells at 6 months; T_14_ = 2.2, P = 0.047). ***C***) There were significantly fewer pyknotic, caspase3^+^ cells in the DG at 6 months compared to 2 months of age (T_13_ = 6.4, P < 0.0001). ***D***) Pyknotic caspase3^+^BrdU^+^ cells were observed at 2 months but not 6 months of age (Mann-Whitney test, P = 0.026). ***E***) Cell distribution within the granule cell layer: box and whisker plots indicate quartiles, plus sign indicates mean. The oldest neurons typically are found in the superficial-most layer (maximum 100) and the youngest neurons are typically found in the deep layers near the hilus (minimum 0). Nearly all immature DCX^+^ cells are found within the deepest 25% of the granule cell layer. P6-born dentate gyrus neurons are distributed throughout, but are most concentrated in the middle of the granule cell layer. There was a significant shift in distribution towards the superficial layers at 6 months of age (T_373_ = 3.2, P = 0.0014). The distribution of DCX^+^ and caspase3^+^ cells did not change between 2 and 6 months of age. While most pyknotic caspase3^+^ cells were found in the deep (neurogenic) layers, dying cells could also be observed in the superficial layers inhabited by P6-born neurons. The location of pyknotic caspase3^+^BrdU^+^ cells are indicated by the filled black symbols. ***F***) Analyses of cells in the superficial layers (superficial 75%). Out of 192 cells examined, only 1 DCX^+^ cell was observed in the superficial granule cell layer, indicating that relatively mature dentate gyrus neurons reside in these layers. The majority of P6-born BrdU^+^ neurons were found in the superficial granule cell layer, with a greater proportion observed at 6 months than at 2 months (T_13_ = 2.7, P = 0.017). At 2 months of age 18% of pyknotic caspase3^+^ cells were observed in the superficial granule cell layer (significantly greater than zero, one sample t-test, T_6_ = 3.2, P = 0.018). The proportion of pyknotic caspase3^+^ cells that were found in the superficial granule cell layer did not differ between 2 and 6 months of age (T_13_ = 0.6, P = 0.5).

To obtain positive evidence for cell death we quantified cells that were both pyknotic and expressed activated caspase3, and found an average of 217 total dying cells per rat at 2 months of age, and 70 cells at 6 months of age (Fig. 4C). Furthermore, we observed on average ~10 pyknotic BrdU^+^caspase3^+^ cells per rat at 2 months of age, indicating death of P6-born cells (Fig. 4D). Since dying BrdU^+^ cells were infrequent and difficult to sample, we also analyzed the anatomical distribution of dying cells within the granule cell layer, where older neurons are located in more superficial regions (Fig. 4E). We first examined the distribution of DCX^+^ cells, since this marker labels immature neurons and can be used to identify regions of the granule cell layer where adult-born cells are dying. As expected, nearly all DCX^+^ cells were located in the deepest 25% of the granule cell layer, near the hilus. P6-born BrdU cells were scattered throughout the granule cell layer and were primarily located in the middle regions. Finally, the majority of pyknotic caspase3^+^ cells were located within the deepest 25% of the granule cell layer, consistent with the death of immature adult-born neurons (Sierra et al., 2010). However, pyknotic caspase3^+^ cells (with and without BrdU) could be found throughout the full width of the granule cell layer at both 2 and 6 months, providing additional evidence that developmentally-born neurons die in young adulthood.

We next quantified the proportions of cells in the superficial 75% of the granule cell layer, i.e. where cells are mature and largely born in early postnatal development (Fig. 4F). As expected, P6-born BrdU^+^ cells were mainly found in this region. However, a significant proportion of pyknotic caspase3^+^ cells were also observed in the superficial granule cell layer region at 2 months (18%; 13% at 6 months, not significantly different) providing additional evidence that death of developmentally-born neurons reflects a modest but significant proportion of overall cell death within the dentate gyrus.

Notably, the distribution of P6-born BrdU^+^ neurons shifted towards more superficial regions of the granule cell layer from 2 to 6 months (Fig. 4E, F). This could reflect ongoing adult neurogenesis in the deeper regions, or preferential death of P6- born neurons in the deeper layers.

## DISCUSSION

Our principal finding is that the survival pattern of DG neurons born in early postnatal development is essentially the opposite of DG neurons born in adulthood (summarized and compared with previously published findings in Fig. 5). In adult rodents approximately 40-80% of adult-born cells die between 1-4 weeks of age (Cameron et al., 1993; Gould et al., 1999; Brandt et al., 2003; Snyder et al., 2009a; Tashiro et al., 2006, McDonald and Wojtowicz, 2005; Mandyam et al., 2007). Remaining neurons continue to survive to at least 6 months in rats (Dayer et al., 2003), 11 months in mice (Kempermann et al., 2003), and likely persist for the life of the animal. In contrast, here we observed no early loss of developmentally-born BrdU^+^ cells between 1-8 weeks but we did find that 17% died after reaching maturity, between 2-6 months of age. A steady loss of P6-born DG neurons between 1-6 months of age has been previously demonstrated in Sprague Dawley rats (Dayer et al., 2003). Here, we confirm these data in another strain (Long Evans), within a more restricted window of development (2-6 months of age), and with specific markers for dying cells. Using behavioral immediate-early gene induction as an indirect measure of circuit integration we found that Fos steadily increased, reaching maximal levels when cells were 4-weeks-old. In contrast, zif268 expression showed a transient peak when cells were 2-weeks-old, suggesting a potential critical period for plasticity in immature DG neurons. While our group sizes were too small for a properly powered comparison between males and females, no obvious sex differences were observed for any measure.

**Figure 5:**
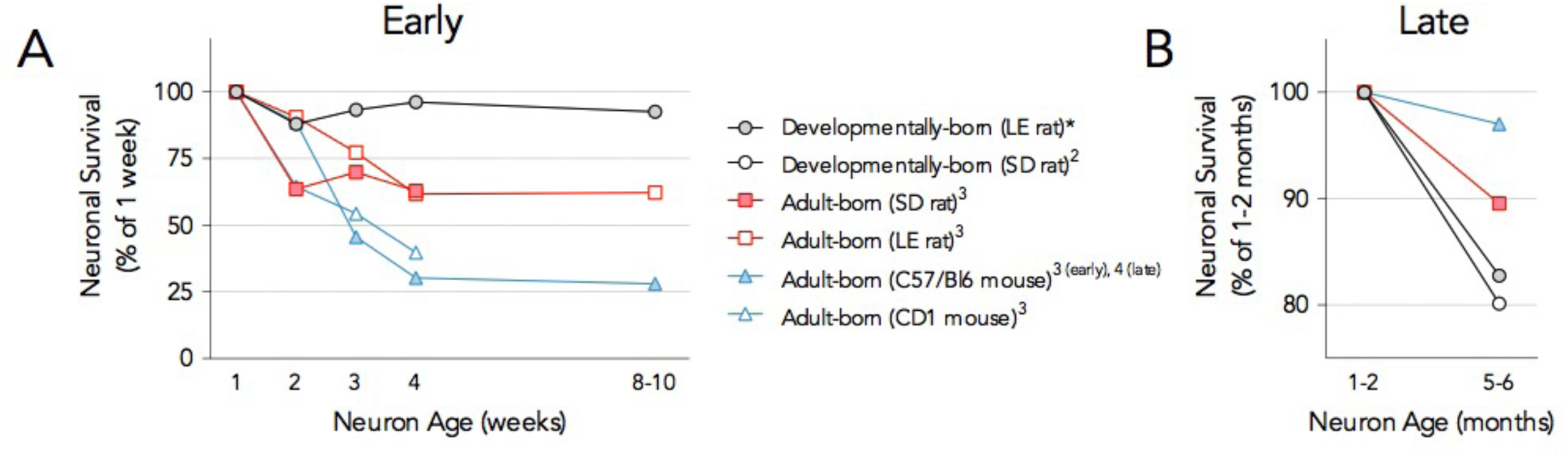
Comparison of early vs. late survival in developmentally-born and adult-born DG neurons. ***A***) Developmentally-born neurons survive early, immature stages of cellular development but many adult-born DG neurons die. ***B***) Roughly 20% of developmentally-born neurons die after reaching maturing but adult-born neurons are relatively stable. *Asterisk indicates data from the current study, other data are from ^2^Dayer et al., (2003), ^3^Snyder et al., (2009a) and ^4^Kempermann et al., (2003). Where raw data were not available, values were extracted from published graphs with Plot Digitizer.

### Alternative explanations for early survival and delayed death

When examining the birth and persistence of cohorts of neurons, it is important to consider possible confounds associated with labeling methods. For example, BrdU that is taken up by precursor cells will continue to label daughter cells with each division until it is diluted below the limits of detection. Redivision of BrdU^+^ cells could therefore give the false impression that P6-labeled neurons are stable between 1-8 weeks after injection, for example if the addition of new cells offset the death of immature cells. Likewise, infrequent/delayed division between 2-6 months could lead to BrdU dilution below the detection threshold, giving the appearance of cell loss. However, these alternative explanations are unlikely to explain our results for several reasons. First, while many BrdU^+^ cells expressed the cell division marker PCNA between 1 hour and 3 days after BrdU injection, when there was corresponding growth in the BrdU^+^ population, there was negligible expression of PCNA from 1 week onwards. Thus, continued division of BrdU^+^ cells cannot explain the early stability (1-8w) or delayed loss (2-6mos) of BrdU^+^ cells. Second, BrdU^+^ adult-born cell numbers remain stable over durations that are significantly longer than those examined here (6-11 months), indicating that incorporated BrdU remains a persistent label throughout the life of postmitotic cells (Dayer et al., 2003; Kempermann et al., 2003). Third, only 5% of 8-week-old cells were non-neuronal (NeuN^−^). Even if this entire population consisted of slowly-dividing stem cells that diluted their BrdU between 2-6 months, this cannot account for the observed 17% drop in P6-born neurons. Fourth, in 2-month-old rats, 18% of dying pyknotic caspase3^+^ cells were located in the superficial granule cell layer, which was devoid of DCX immunoreactivity, strongly suggesting that mature neurons were actively undergoing apoptosis. Fifth, the presence of pyknotic caspase3^+^BrdU^+^ cells provides direct evidence that P6-born neurons undergo apoptosis in young adult rats. Assuming a linear decline in cell death from 10 BrdU^+^ cells at 2 months to 0 BrdU^+^ cells at 6 months (5 cells on average) and 1 hour to clear apoptotic DG neurons (Sierra et al., 2010): 5 dying BrdU^+^ cells x 2880 hours = 14,400 cells predicted to die between 2-6 months (we observed 28,538 BrdU^+^ cells lost). Sampling limitations and insufficient knowledge about the kinetics of cell clearance make precise calculations difficult, but the BrdU and pyknotic/caspase data are broadly consistent and indicate that substantial numbers of developmentally-born neurons die throughout young adulthood.

### Maturation and early zif268 expression relative to adult-born dentate gyrus neurons

Morphological and electrophysiological studies have found that developmentally-born neurons mature faster than adult-born neurons (Overstreet-Wadiche et al., 2006; Zhao et al., 2006). While expression patterns of DCX, NeuN and Fos did not suggest obviously different maturation rates compared to adult-born neurons, the zif268 profile was shifted 1 week earlier than we have previously observed in adult-born neurons. Specifically, we have previously observed a sharp peak in zif268 expression in 3-week-old adult-born neurons (Snyder et al., 2009a); here we observed a similar peak in 2-week-old developmentally-born DG neurons. This peak was not observed in the overall DG population, indicating that it is related to cell age rather than the developmental stage of the animal. Zif268 is an immediate-early gene that is critical for long-term plasticity and long-term memory (Jones et al., 2001). Moreover, zif268 promotes the survival and experience-dependent activation of immature adult-born neurons (Veyrac et al., 2013). If zif268 plays a similar role in developmentally-born neurons, our findings suggest that neurogenesis around the peak of postnatal DG development (~P6) would result in a very large population of highly plastic neurons 2 weeks later when rats are ~3 weeks old and just beginning to display hippocampal-dependent memory (Rudy, 1993; Akers and Hamilton, 2007; Raineki et al., 2010). While the immediate-early genes zif268, Fos and Arc are often used as cellular activity markers, few studies have compared patterns of expression. All 3 immediate early genes are upregulated by spatial water maze training, but Arc has been found to correlate best with performance and task demands (Guzowski et al., 2001). Arc and Fos are largely expressed by similar DG cell populations (Stone et al., 2011) but here we find that Fos and zif268 are only partially co-expressed within cells. While most Fos^+^ cells also expressed zif268, zif268 was present in a much larger population of cells that did not express Fos, possibly because zif268 has a lower threshold for activity-dependent expression (Worley et al., 1993; 1991). It is worth noting that since we did not include an unstimulated, caged control group, we cannot conclude with certainty that zif268 and Fos were induced by novel context exposure, particularly in light of evidence that immature adult-born DG neurons express high levels of zif268 in the home cage (Snyder et al., 2012b; Huckleberry et al., 2015). However, the expression timecourse is consistent with the formation of synapses, and others have shown that resting levels of immediate early genes are activity-dependent (Worley et al., 1991) and experience-specific (Marrone et al., 2008). We therefore believe that these markers are valid activity indicators, but future studies are required to elucidate their precise functions.

### Neuronal persistence, turnover and memory

The initial persistence and delayed death of developmentally-born DG neurons raises fundamental questions about their role in hippocampal function. Clearly, the early postnatal period is experientially rich; high mnemonic demands may require the complete survival of developmentally-born neurons during their first few weeks and months. That experience can increase the survival of adult-born neurons is consistent with this idea (Kempermann et al., 1997a; Gould et al., 1999; Epp et al., 2007, Tashiro et al., 2007), but the extent to which developmentally-born neurons are hard wired for survival, or survive based on sensory experience and mnemonic demands is not clear. Additional investigation into the early stability of developmentally-born neurons is therefore warranted, particularly in light of intriguing evidence that behavioral stimuli can induce death of superficially-located (presumably developmentally-born) granule cells (Olariu et al., 2005).

In our late survival experiment, P6-born DG neurons died between 2 and 6 months of (cell) age. While death of mature neurons is observed in neurodegenerative disorders, it is generally not believed to occur in the healthy young adult brain. Our findings suggest that death of mature functional DG neurons may be a part of normal development/aging. The consequences of dying developmentally-born neurons would be very distinct from death of immature adult-born cells, as they would have presumably participated in memory processes and their removal could result in loss of information from hippocampal circuits. In contrast, dying immature adult-born cells would have had fewer, if any, opportunities to store memories or process information prior to their death.

Computational models predict that adult neurogenesis coupled with neuronal turnover benefits learning (Becker, 2005) beyond what can be achieved with adult neurogenesis alone (Meltzer et al., 2005). The addition of new neurons and removal of old neurons might preferentially enhance new learning at the expense of retaining older information (Meltzer et al., 2005; Chambers and Conroy, 2007). Indeed, there is evidence to support these predictions: in songbirds, the death of mature neurons promotes the birth of new adult-born neurons, and neuronal turnover contributes seasonal changes in song repertoire (Alvarez-Buylla and Kirn, 1997; Larson et al., 2014). In mammals, adult neurogenesis causes forgetting of hippocampal memories (Akers et al., 2014) and consolidation of memory into extra-hippocampal structures (Kitamura et al., 2009). Our data raise the question of whether death of developmentally-born neurons might also contribute to the loss and consolidation of hippocampal memory.

While the number of dying developmentally-born cells is a considerable proportion of the total DG population, the majority of P6-born DG neurons did not die between 2-6 months of age. This fits with behavioral findings that, while some memories may transform or be forgotten over time, episodic-like memories can persist indefinitely in the hippocampus (Moscovitch et al., 2016). The combination of survival and death could therefore be highly adaptive, enabling detailed memory for important events while minimizing overreliance on obsolete information (Richards and Frankland, 2017).

### Relevance for mental health

Animal models indicate that genetic factors and experience in both early life and adulthood can impact total DG cell number (Kempermann et al., 1997b; 1997a; Fabricius et al., 2008; Oomen et al., 2011). Our findings are therefore relevant for a number of psychiatric disorders. For example, the restricted window of zif268 expression might render specific cohorts of neurons vulnerable to neurodevelopmental insults to the hippocampus (Alberini and Travaglia, 2017). Altered production and survival of developmentally-born DG neurons is also relevant for a number of disorders that are associated with large-scale structural changes in the hippocampus, and sometimes the DG in particular, such as autism (Saitoh et al., 2001; Schumann et al., 2004), schizophrenia (Tamminga et al., 2010; Lodge and Grace, 2011) and depression (McKinnon et al., 2009). For example, recent reports indicate that depressed patients have fewer total DG neurons in the anterior hippocampus, which can be restored by SSRI antidepressants (Boldrini et al., 2013; 2014). While antidepressant treatments are often cited for their proneurogenic properties, our findings indicate that changes in the developmentally-born cell population could also contribute to changes in total granule cell number.

Whether it is in fact desirable to rescue developmentally-born neurons is another question, as they appear to be fundamentally different from (even relatively old) adult-born neurons. Adult-born neurons have greater experience-dependent morphological plasticity (Tronel et al., 2010; Lemaire et al., 2012), display unique experience-dependent patterns of immediate-early gene expression (Snyder et al., 2009c; Snyder et al., 2011; Tronel et al., 2014), and have distinct functions in contextual encoding (Nakashiba et al., 2012; Danielson et al., 2016). Thus, culling developmentally-born neurons and replacing them with new neurons may in fact be beneficial, particularly if the developmentally-born neurons are less plastic and have formed maladaptive associations.

## Conclusions

Lifelong neurogenesis in the DG results in a degree of cellular heterogeneity that has only begun to be explored. Here, we examined how cells born at a single time point survive from infancy through young adulthood. How do our results generalize to neurons born at, and surviving to, other stages of development? Assuming there is delayed death of other DG neurons born in the first postnatal week, there could be hundreds of thousands of cells lost in young adulthood in rats, and perhaps even more in older age. The survival and death of different cohorts of adult-born neurons are interrelated processes (Dupret et al., 2007) and it is known that mature DG neurons regulate the recruitment of adult-born DG neurons (Alvarez et al., 2016; McAvoy et al., 2016). Thus, there may be a functional link between the loss of developmentally-born neurons and the addition of adult-born neurons. Identifying such relationships may help resolve longstanding conflicting reports that DG neurons accumulate (Bayer, 1982a; Bayer et al., 1982b; Amrein et al., 2004), remain constant (Boss et al., 1985; Rapp and Gallagher, 1996; Amrein et al., 2004) or fluctuate throughout the lifespan (Boss et al., 1985).

## Acknowledgements

This work was supported by an NSERC Discovery Grant (JSS), an NSERC postgraduate scholarship and Killam Doctoral Scholarship (SPC).

